# Annotation-free discovery of functional groups in microbial communities

**DOI:** 10.1101/2022.08.02.502537

**Authors:** Xiaoyu Shan, Akshit Goyal, Rachel Gregor, Otto X. Cordero

## Abstract

Recent studies have shown that microbial communities are composed of groups of functionally cohesive taxa, whose abundance is more stable and better associated with metabolic fluxes than that of any individual taxon. However, identifying these functional groups in a manner that is independent from error-prone functional gene annotations remains a major open problem. Here, we develop a novel approach that identifies functional groups of taxa in an unsupervised manner, solely based on the patterns of statistical variation in species abundance and environmental parameters. We demonstrate the power of this approach on three distinct data sets. On data of replicate microcosm with heterotrophic soil bacteria, our unsupervised algorithm recovered experimentally validated functional groups that divide metabolic labor and remain stable despite large variation in species composition. When leveraged against the ocean microbiome data, our approach discovered a functional group that combines aerobic and anaerobic ammonia oxidizers, whose summed abundance tracks closely with nitrate concentrations in the water column. Finally, we show that our framework can enable the detection of species groups that are likely responsible for the production or consumption of metabolites abundant in animal gut microbiomes, serving as a hypothesis generating tool for mechanistic studies. Overall, this work advances our understanding of structure-function relationships in complex microbiomes and provides a powerful approach to discover functional groups in an objective and systematic manner.

## Introduction

Microbial communities often involve thousands of different taxa (e.g., 16S rRNA ribotypes, strains, etc.) despite their sharing of many metabolic functions^1,2^. This observation has led to the notion that the major metabolic fluxes in a community of microbes are better captured not by the overall taxa composition, but by the abundance of a much smaller set of so-called ‘functional groups’ or guilds^3–5^, akin to those defined for plant and animal communities^6–8^. Members of the same group can fluctuate widely in abundance, replacing each other across space and time due to (often stochastic) ecological interactions, like predation, which can also be highly strain specific^9,10^. In contrast, the abundance of the group as a whole and its combined metabolic output – i.e. the ecosystem service provided by the group, remains stable and is better-associated with the environmental parameters (pH, nutrient concentration, etc.) that control metabolism^3,5^. Thus, as in the case of financial portfolios in which diversification stabilizes returns^11^, functional groups buffer microbial ecosystems from the intrinsic volatility of species dynamics. Moreover, functional groups also provide a more interpretable and simpler description of microbial ecosystems – in terms of key metabolic functions and the groups performing them – than the more readily accessible taxa and gene catalogs.

Despite consensus about the importance of functional groups in microbial ecology, identifying and discovering them is still a major challenge. This is a problem that is not specific to microbial ecosystems. For animals and plants, functional groups are defined based on shared traits, but more often than not it is unclear what the relevant traits are and how to measure them in a systematic manner^12^. As a result, there can be a large degree of subjectivity in how functional groups are defined, compromising the value of the approach. Traits such as manner of feeding for insects are much harder to define for microbes, and the best analog we have is phenotypic inference based on genome annotation of metabolic capabilities or taxonomy. However, these annotations carry a high degree of uncertainty^13–15^, especially when considered in the context of a community. At the core of the problem is that thorough phenotypic characterizations are rarely available for microbes, even for the few that have been cultured. As a consequence, functional gene annotations rely on what is known for a handful of model organisms, like *E. coli* and *B. subtilis*. The less related the organism of interest is to these well-studied model systems, the lower the quality of annotations^16^. To complicate matters further, the presence of a pathway in a genome, even if uncontroversial, does not imply that it is utilized by the organism under the conditions in which the community exists^17,18^. Therefore, the viability of annotation-based approaches is limited to special cases in which there is a clear, unambiguous relation between taxa identity and function, e.g., oxygenic photosynthesis and cyanobacteria^3^.

To address this challenge, here we present an unsupervised, annotation-free approach that identifies functional groups of taxa based on the patterns of statistical variation in microbiome composition and environmental variables. The method identifies the minimal group of taxa that best correlate to the levels of a given environmental parameter, like the abundance of a metabolite or the concentration of a nutrient. If the environment is constant across samples, the approach we present in this paper can identify groups of taxa that are most stably maintained across replicates, likely representing ‘species portfolios’ with redundant function. Below we provide a detailed explanation of the approach and show how it can successfully be applied across various data sets, ranging from replicate microcosms^4^ to the TARA Oceans microbiome^19^ to metabolomic profiles of animal gut microbiomes^20^.

## Results

### Theory: Extracting functional groups from microbiome data

We interpret the challenge of finding functional groups as an optimization problem, i.e. finding the smallest group of taxa that maximize the correlation between their combined abundance and a given environmental variable. To gain intuition into how to solve this problem, consider a group of two taxa and one variable, *y*, representing, for instance, the concentration of a metabolite. From a statistical standpoint, what would make the group a better predictor of *y* than individual taxa (Figure 1A-B)? The answer depends on how the two species covary with *y*, as well as with each other. It is straightforward to show that, for the coefficient of determination to increase when the two taxa are counted as one, their individual abundances should not be strongly positively correlated (Figure 1C-D). By contrast, if the two species are anticorrelated or uncorrelated in abundance, they may complement each other – the species may be good predictors of *y* in non-overlapping subsets of samples, thus increasing the overall value of the correlation between the group and *y*. At the same time, if one species has a positive covariance with y and the other a negative covariance, their individual effects may cancel out. As shown below, these intuitions can be generalized: the best functional group is one in which members’ abundances tend to be correlated with the external variable in a consistent manner (same sign), while at the same time complementing each other (uncorrelated or anticorrelated). Solving this tension between consistency and complementarity is the optimization challenge.

**Figure 1.**
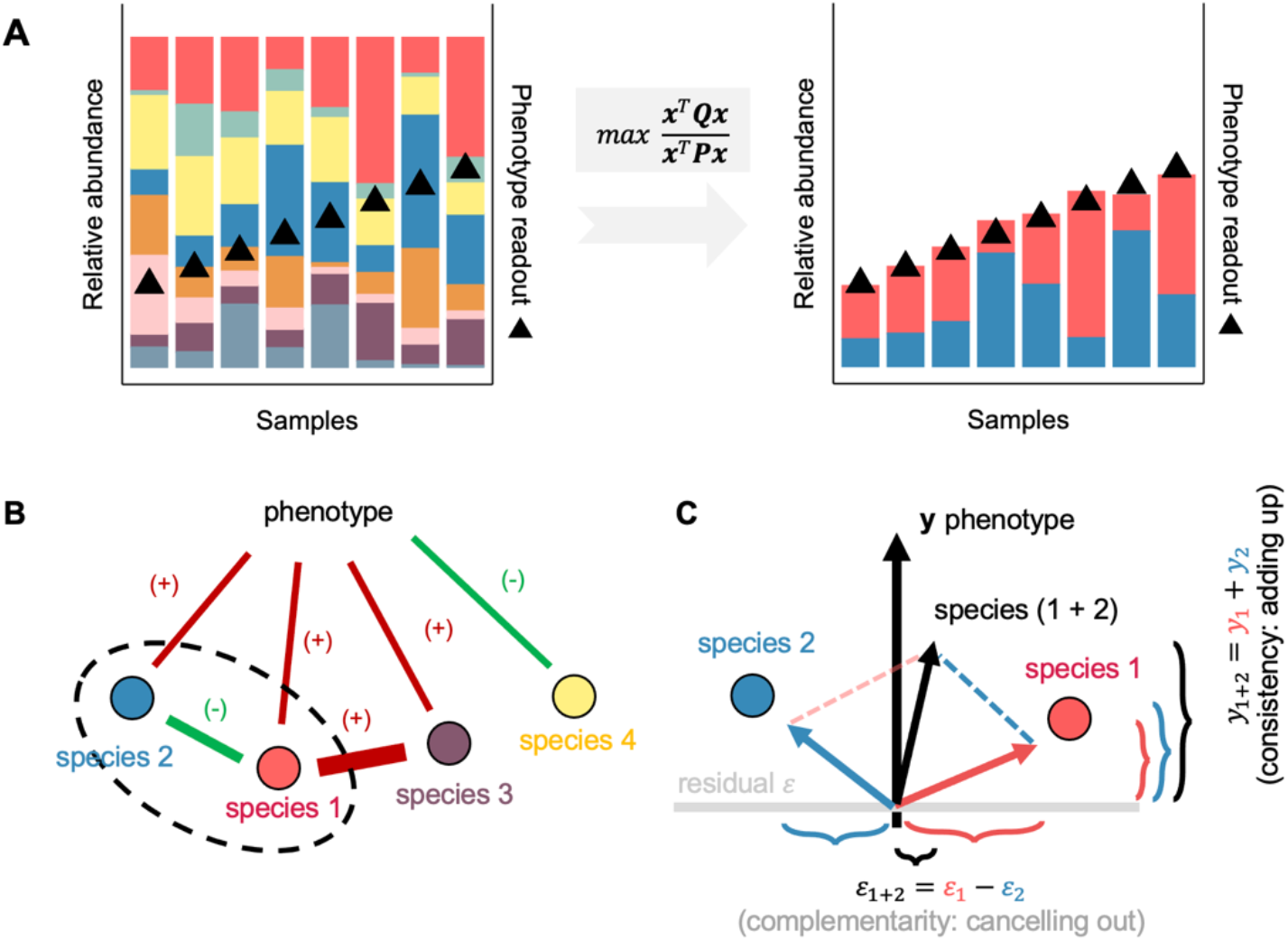
Schematic illustration of Ensemble Quotient Optimization (EQO). (A) A typical microbiome is composed of a diversity of taxa, of which individual taxa are only poorly coupled with the measured phenotypic readout. (B) With EQO, species 1 (red) species 2 (blue) are selected to be grouped into an assemblage, whose relative abundance is strongly correlated with the phenotypic readout. Species 1 and species 2 are both positively correlated, though not strongly, with the phenotype (r_1,y_ = 0.41, r_2,y_ = 0.53), while they are anti-correlated with each other (r_1,2_ = - 0.54). They are both *consistent* and *complementary*. (C) Consistency implies that two species “add up” in the direction of the phenotype axis while complementarity implies that two species “cancel out” each other with their residuals orthogonal to the phenotype axis. Black vertical axis marks the phenotypic variable. Gray horizontal axis marks the residual of species after projecting onto the phenotypic variable axis.

We derived a simple mathematical expression that captures this tension and that can be used to find functional groups of taxa, as here defined. Consider a community matrix with the abundances of *n* species abundances over *m* samples, and an environmental variable *y*. A group of taxa is defined by a vector of ***x*** ∈ (0,1)^*n*^, where the *i*^th^ position is 1 if the corresponding species belongs to group, or 0 otherwise. The correlation between the group abundance and *y* can be expressed in a compact form, which we term the Ensemble Quotient, 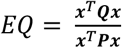, where *P* and *Q* are algebraic transformation of the community matrix that capture the covariance between species (*complementarity*), and the covariance between species and *y* (*consistency*), respectively (Methods and Supplementary Notes). The exact form of these algebraic transformations depends on whether *y* is a continuous variable (e.g. a metabolite concentration), a categorical variable (e.g. healthy or disease states) or a constant (in cases when we want to identify stable groups from biological replicates) (Figure S1). Independently of the type of variable *y* represents, the *EQ* is the objective function we want to maximize over ***x***.

Two problems that arise when searching for a group that maximizes *EQ* are the large number of possible solutions and the risk of overfitting. With only 100 species, there are over 75 million groups of size 5, and this number increases exponentially with species richness. As described in Methods, we circumvent this problem using a genetic algorithm that allows us to efficiently search functional groups in communities with hundreds of taxa. The risk of overfitting appears because, the larger the group, the easier it should be to find a species combination that produces a good fit. We solve this problem by putting a penalty on group size (regularization) when appropriate and by assessing the statistical significance of out-of-bag predictions.

We named the functional group discovery approach EQO, for Ensemble Quotient Optimization. Below, we illustrate its power on three distinct datasets: a set of communities assembled through controlled laboratory microcosms^4^ in which amplicon sequence variants (ASVs) fall into two, phylogenetically constrained, functional groups; the TARA oceans microbiome data^19^ coupled with environmental parameters such as the nitrate or oxygen concentration in the water column, and an animal microbiome dataset^20^ in which taxa composition data is paired with detailed metabolomic profiling.

### Stable functional groups in replicate microcosms

We start by applying EQO to a data set developed from laboratory scale enrichments of soil communities under controlled conditions. The communities were assembled by serial passaging in minimal media with glucose as the limiting resource until they reached a stable composition^4^. Despite identical environmental conditions among replicates, at the fine-grained level of genetic resolution (ASVs), replicate microcosms stabilized into communities with very different compositions. However, the communities were much more similar to each other at the level of major taxonomic families, with two families (*Enterobacterales* and *Pseudomonadales*) dominating the assemblage. It was later confirmed that these two families constituted bona fide functional groups, with one group (*Enterobacterales*) performing partial glycolysis, and the other (*Pseudomonadales*) performing gluconeogenesis to complete the full respiration of carbon. Because this is a small scale, controlled experiment, with experimentally validated functional groups, it serves an ideal case to test whether out approach can reproduce results based on taxonomic annotations.

Importantly, this is a case in which we do not have an external signal to guide the functional group search, since all replicate communities were assembled from the same inoculum and under the same environmental conditions. Instead, we want to find the partitioning of the community that is most stable across replicates. To this end, we define y, the environmental variable, as a constant across all replicate microcosms. By finding the group of taxa that best correlates with this constant value, we identify the most stable bi-partition of the community. The EQ formulation for a constant *y* is described in Methods.

Remarkably, our unsupervised approach was able to reproduce the bipartition of the community into *Enterobacterales* and *Pseudomonadales*, but without any prior information about taxonomy, purely from the statistical patterns of ASV variation in the data (Figure 2, 94.0% ± 2.2% of the reads correctly mapped; Pearson’s r = 0.97 between the supervised and unsupervised groups). Interestingly, in this specific case we did not need to penalize group size. The optimal group, without constraining for size, is the one that partitions the community into the two, experimentally validated functional groups. Overall, these results show that the approach here developed is capable of identifying functional groups. Our next step is to test its value on real world, environmental data.

**Figure 2.**
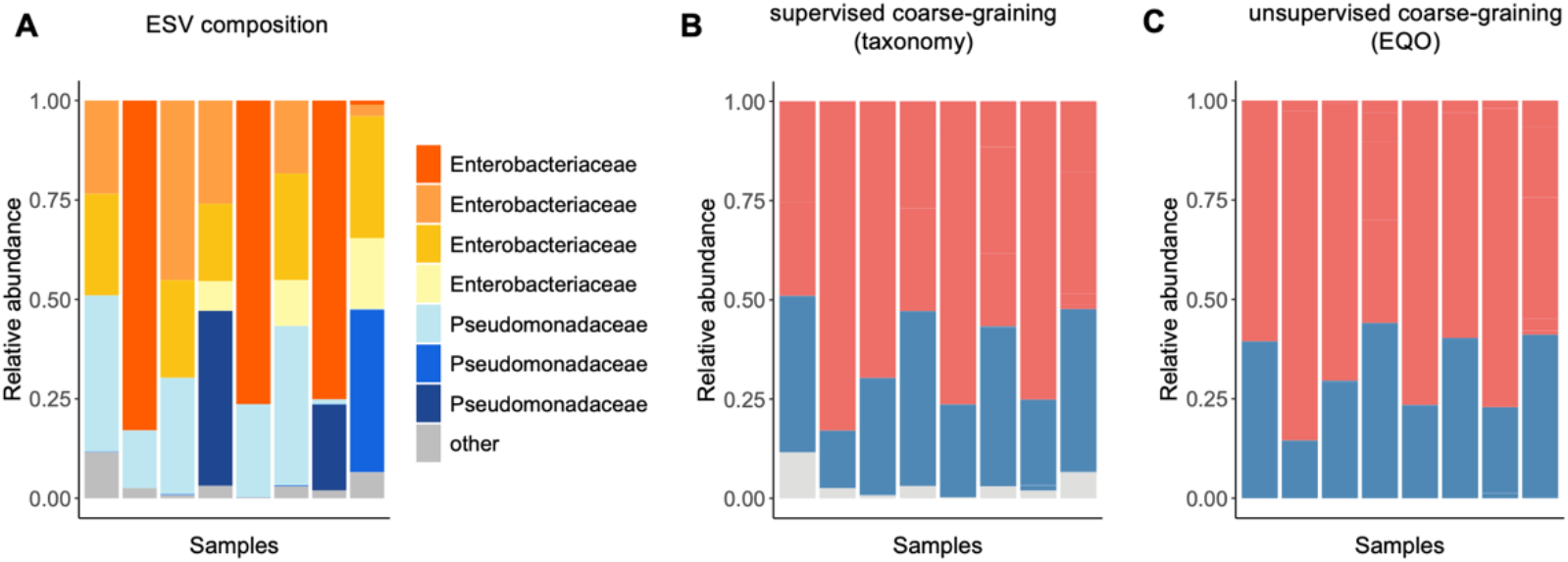
Coarse-graining functional groups in replicate microcosms. (A) original ESV-level composition of the replicate microcosms assembled with glucose as carbon source. (B) Supervised coarse-graining into Family-level groups based on taxonomic annotation. (C) Unsupervised coarse-graining by EQO for groups with stable relative abundances across samples.

### Nitrogen cycling in the ocean microbiome

To study EQO’s performance in a complex, environmental datasets, we applied it on the TARA oceans dataset^3,19^. Samples in this data set originate from depths ranging from 5.3 to 792 meters and contain a total of 2451 taxa classified at the genus level. Of those, 97 are over 1% abundance in at least one sample. Besides depth, other environmental variables are pH, temperature, nitrate, phosphate, silicate and oxygen, with most of them systematically changing as a function of depth. Of these, we chose to focus on nitrate because of its importance in nutrient cycling, and because it is a direct intermediate of microbial metabolism – a product of nitrification and a substrate for denitrification.

EQO was able to discover a functional group of genera involved in nitrogen cycling in the ocean. Using the Akaike Information Criterion (AIC), calculated as –2*k* - ln(*L_k_*), where *L_k_*, is the regression likelihood and *k* the group size, we established an optimal group size of 11 members (Figure S2). To estimate the relative contribution of each individual member, or member-member pairs to the group’s performance, we took advantage of the large number of samples in TARA to apply cross-validation (Figure 3A). We perform cross-validation by dividing the data in training and test (50-50) sets and iterating 1000 times, resulting in an equal number of potentially different groups. We estimated the relative importance of a group member, *i*, by summing the R^2^ of all groups in which *i* was present (∑ *x_i_R*^2^), and similarly for pairs of members (∑ *x_i_x_j_R*^2^).

**Figure 3.**
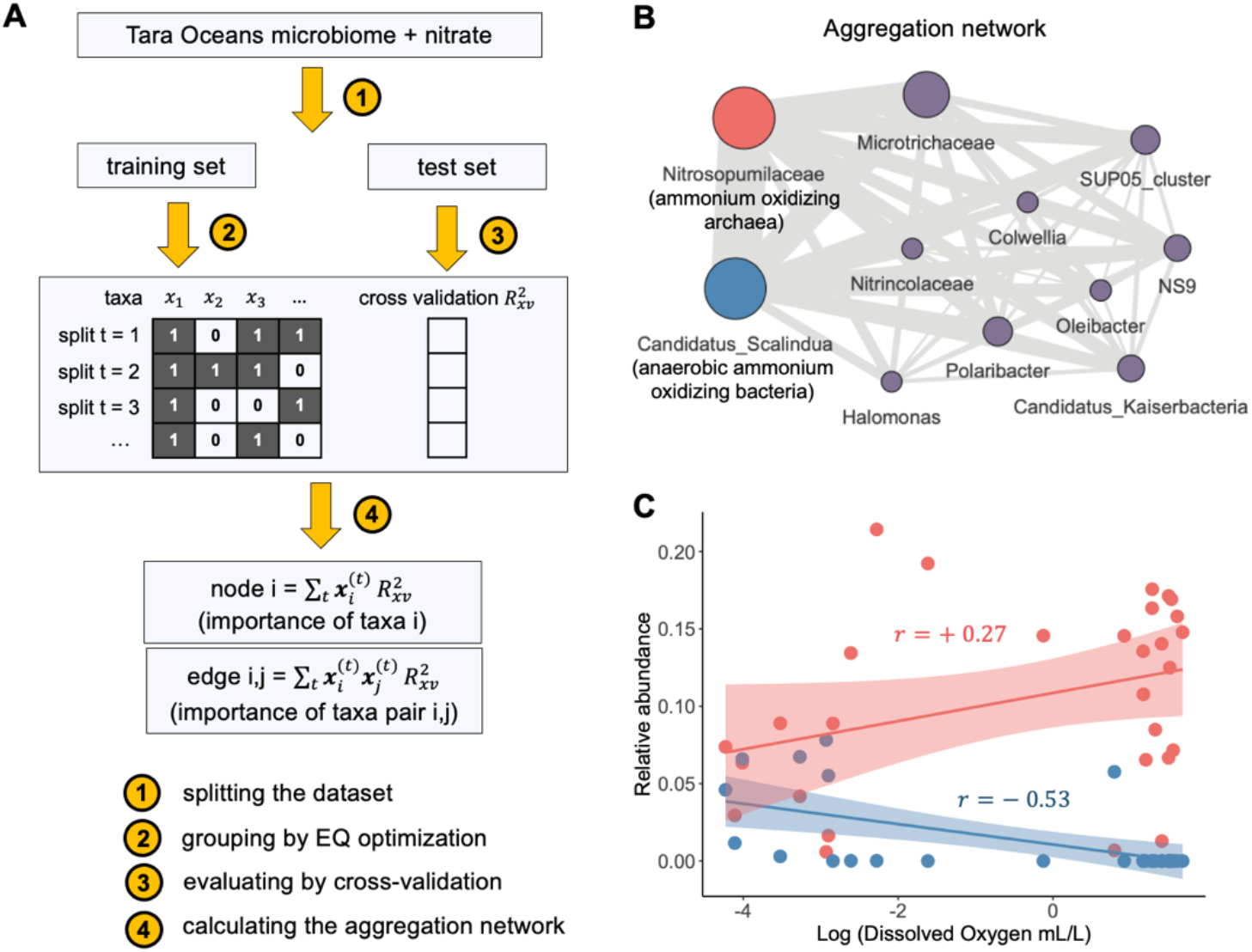
Functional guilds of nitrogen cycling in the ocean microbiome. (A) A cross-validation-based algorithm to construct the aggregation network for functional grouping (Methods). The microbiome dataset with the accompanying metadata is randomly split into a training set and a test set. The EQO was applied to the training set to generate a best assemblage, which was then validated with the test set for cross-validated R^2^. After iterating 100 times with the cross-validation process, the cumulative cross-validated R^2^ for assemblages where a single taxon was present or a pair of taxa was co-present were calculated to indicate the node size and edge width in the aggregation network. (B) Aggregation network for nitrate concentration of the Tara Oceans microbiome. *Nitrosopumilaceae*, the ammonium oxidizing archaea (highlighted in red) and Candidatus *Scalindua*, the anaerobic ammonium oxidizing bacteria (highlighted in blue) tended to be always co-selected by the algorithm, which led to strong cross-validated R^2^ with nitrate concentration, suggesting these two taxa together formed a functional guild. (C) *Nitrosopumilaceae* and Candidatus *Scalindua* showed opposite trend of variation in response to dissolved oxygen in the water column, indicating that they are alternating based on the level of oxygen availability.

Out of the 11 members, two taxa, *Nitrosopumilaceae* and Candidatus *Scalindua*, had the highest relative contribution to group’s ability to predict nitrate concentrations. *Nitrosopumilaceae* is a well-known clade of ammonia oxidizing archaea^21^, while Candidatus *Scalindua* is the most abundant anaerobic ammonia oxidizing (annamox) bacteria in marine environment^22^. These two taxa are always co-selected by EQO during cross-validations, as shown in the network of relative pair importance (Figure 3B). This is because these two taxa alternate across sampling stations as a function of oxygen concentration (across samples where both taxa are >1% relative abundance, their correlation is Pearson’s r = - 0.64), in complete agreement with their predicted roles as aerobic and anaerobic ammonia oxidizers (Figure 3C). Because conditions with low oxygen concentration are rare in the TARA samples, the correlation between Candidatus *Scalindua* alone and nitrate is weak (r = 0.31). However, when combined with *Nitrosopumilaceae*, these two taxa complement each other and the correlation was enhanced to r = 0.82 (r = 0.89 for the whole group of 11 taxa). These results show that nitrate concentration in the water column is tightly controlled by the abundance of ammonia oxidizing *archaea* and bacteria, to the extent that nitrate measurements are a good proxy for the overall abundance of this functional group.

### Mapping metabolites to functional groups in animal microbiomes

One of the most compelling potential applications of EQO is the identification of functional groups responsible for the production or consumption of metabolites in gut microbiome data. We leveraged an animal gut microbiome dataset with 101 fecal samples from a wide range of 25 mammalian species, accompanied by 74 peak features that are detected by GC-MS in >80% samples^20^. Unlike the case of nitrate discussed above, which has been a major focus of research in since the early days of environmental microbiology, the metabolic processes that drive the production or consumption of metabolites in animal guts are much less understood. This makes interpreting the results of the algorithm much harder and demands that we focus only on those predictions that are well-above any significance threshold.

We took a two-fold approach to make sure we focus on statistically significant functional group predictions (Figure S3). As before, we use cross-validation to assess out-of-bag predictions. Of all the AIC-based optimal groups, 77% of groups with cross-validated *R^2^* (xv*R^2^*) below 20% were filtered from further analysis (e.g., most small groups (1-3 taxa) have xv*R^2^* ~ 0). Groups with xv*R^2^* > 20% were composed of 7 ± 1 taxa. For these groups, we asked whether their xv*R^2^* was statistically significant relative to random groups of the same size. We used an p-value adjustment, multiplying the p-values by a factor 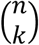, where *n* is the total number of taxa in the microbiome and *k* is the number of taxa in the group, to account for the fact that the number of hypotheses to test increases rapidly with group size. After accepting groups with adjusted p-values below 0.01, we ended with 12 (16.2%) metabolites with a significant functional group prediction (Figure 4A).

**Figure 4.**
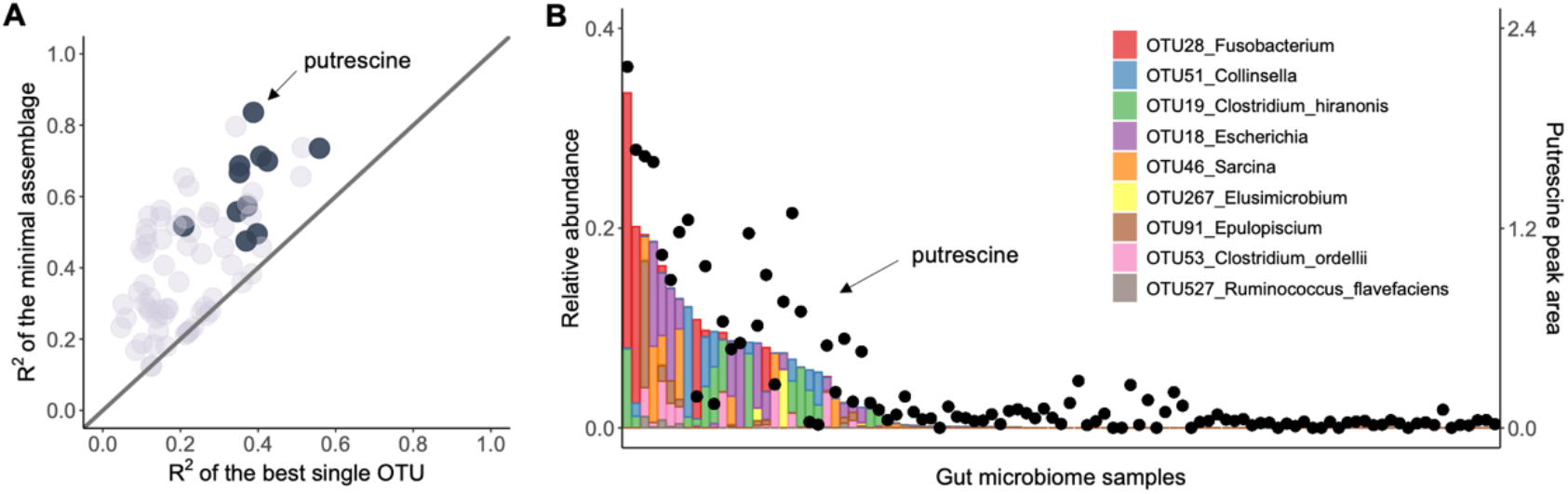
Predicting the level of metabolites with minimal assemblages in gut microbiome. (A) Linear regression R^2^ of the best single OTU compared to the linear regression R^2^ of the best minimal assemblages for 74 metabolites prevalent in > 80% samples. Metabolites that passed significance tests were highlighted as dark black dots (Methods). (B) Putrescine is among the metabolites that passed the stringent significance test, whose level can be strongly predicted by a minimal assemblage of 9 OTUs, which are alternating across different host animals. Host animals are ranked based on descending order of putrescine detected. Full details of animal hosts are shown in Figure S4.

The best predicted metabolite was putrescine (1,4-di-amino-butane), a common polyamine molecule derived from amino acids such as arginine^23^. Polyamines such as spermidine and putrescine have been found with remarkable importance in aging^24^, cognitive function^25^, inflammation suppression^26^ and cancer development^27^. Putrescine is mainly enriched in animals such as leopard, lion and tiger, bear and coati, compared to those herbivores such as elephant, rhino and zebra, consistent with the idea that it is a derivative protein metabolism (Figure 4B and Figure S4). Of the different group members selected by EQO, an operational taxonomic unit (defined at a 97% cutoff) of the genus *Fusobacteria* had the highest relative importance (Figure 4B). *Fusobacteria* have been experimentally found to be enriched in animal systems with high production of putrescine and is known to be able to synthesize it both in vitro and in vivo^28,29^. Other taxa in the group such as *Clostridium* might also play a role in putrescine production through anaerobic fermentation of proteins. Given the diversity of polyamine biosynthesis pathways and microbes that carry them, the production of putrescine is a good example of a function that is better understood as the result of the collective action of multiple species^30,31^. Our framework allows us to deal with this scenario and generates hypotheses regarding the processes and players involved in polyamine production.

## Discussion

Over the last couple of decades, DNA sequencing technologies revolutionized the study of microbial communities, by allowing us to construct catalogs of genes and taxonomic markers in a rapid, high-throughput and inexpensive manner. However, rarely are species and gene catalogs themselves the direct subject of a research question, but rather the variables at our disposal to address questions that relate to what functions microbes mediate: anaerobic respiration, digesting organic matter, producing a toxin, etc., are examples of the functions that microbes mediate. Finding a way to map from the variables we can measure (taxa and genes) to the ones we care about (function), or in other words, solving the structure-function mapping, is a primary intellectual challenge in the field of microbial ecology. Current approaches are bottom-up in nature, building a picture of the community based on functional gene annotations or taxonomic descriptions. We argued that this approach is limited. Instead, here we proposed a framework to coarse-grain communities into functional groups in an unsupervised manner by exploiting the statistical patterns of taxonomic abundance microbial communities and the metadata associated with it. We showed that this approach is surprisingly effective at identifying functional groups without the need for functional or taxonomic annotation in synthetic and natural datasets. The apparent success of EQO suggests that high-throughput surveys of microbial environments in which microbial composition data is paired with metadata (e.g. nutrient levels) can be systematically leveraged to study the structure-function mapping of microbial ecosystems.

EQO builds on the assumption that members of a functional group covary weakly or negatively with one another – a basic tenet of portfolio theory^11^. However, this is not always the case. During early stages of community assembly, e.g. during the colonization of a new environment, environmental filters couple the dynamics of functionally equivalent species^32^. In contrast, as community composition approaches a steady state, functional group members can be decoupled by various factors, including historical contingencies, different sensitivities to environmental parameters (O2 concentration), and biotic interactions, to name a few^33^. It is in this scenario, where the community composition is not undergoing early successional changes, that approaches such as ours can be applied.

Microbial communities can be composed of many thousands of taxa, especially if defined at high levels of genetic resolution, like strains. If diversity is too high (e.g., 1000 taxa or more) the unsupervised identification of functional groups may become too impractical due the explosion of the search space size. However, as shown in this paper, even in systems with high natural diversity, like oceans or gut microbiomes, we have been able to constrain the search space to less than 100 taxa. This reduction in complexity relied on two inherent aspects of microbiome data. The first one is that rank abundance curves have very long tails, meaning that although total species counts are large (tens of thousands), the vast majority of taxa appear in very few (e.g., only one) samples. Since those extremely rare taxa do not carry any useful information in the statistical sense, we discard them from the analysis, drastically reducing search space dimensions. Second, taxa can be collapsed at the different levels of phylogenetic resolution (strains, species, genera, families, etc.). For instance, with the Tara Oceans dataset we chose to collapse taxa at the level of genera (e.g., 97% similarity cutoff in 16S rRNA), a common practice in the field, before applying EQO. Although this choice proved effective, it is partly ad-hoc and more systematic approaches to reduce phylogenetic dimensions should be explored.

We have presented the first systematic and unsupervised approach to resolve the structure function mapping of microbial communities. The approach relies on study designs that measure microbiome composition and environmental measurements across multiple sampling stations (ideally hundred or more). Yet, most metagenomic datasets typically lack this structure, either because of the low number of samples or the lack of functional measurements. As we continue to develop the approach, we hope our work clarifies the critical importance of designing microbiome surveys in suitable for the discovery of structure-function relations.

## Methods

### Microbiome datasets

Analyses of replicate microbial microcosms^4^ and gut microbial communities^20^ were based on Amplicon Sequence Variants (ASVs) of 16S rRNA gene amplicon sequencing generated by the original authors. Analysis on the Tara Oceans microbiome was based on closed-reference taxonomic clustering of metagenome-extracted 16S miTAGs^3,19^. Briefly, 16S rRNA short reads extracted from metagenomes by the original authors were mapped to SILVA 138 reference database^34,35^ at 99% similarity with vsearch v2.21^36^. Considering the large number of sequence clusters (~ 40,000) generated from short reads (~100 bp), we used taxonomic composition resolved at the genus level as the input for our algorithm. Reads that are unable to be classified at the genus level are binned to the finest classifiable taxonomic level. Environmental metadata for the Tara Oceans microbiome including nitrate concentration and dissolved oxygen level as well as GC-MS measurement of metabolites for the animal gut microbiome were requested from the original authors of the previous studies^3,20^.

### Formulation of the Ensemble Quotient

Microbiome coarse-graining towards uniform, continuous or categorical phenotypic variables can be generalized into an optimization problem as follows

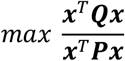

where ***x*** is a Boolean vector of length *n* to be solved from the optimization, for which 1 or 0 represent presence or absence of a species in the ensemble. The number *n* indicates the dimension of the microbiome, e.g., the number of species in an OTU table. For instance, ***x*** = [1,1,0,1,0] indicates that there are in total 5 species in the microbiome, in which the 1st, 2nd and 4th species should be coarse-grained into the ensemble. Matrices ***P*** and ***Q*** are given by the algebraic transformation of the taxa and phenotype, with slightly different forms for uniform, continuous or categorical phenotypic variables detailed as follows. Derivation of the whole mathematical framework is detailed in Supplementary Notes.

a. uniform phenotypic variable (i.e., composition of the assemblage is stable)

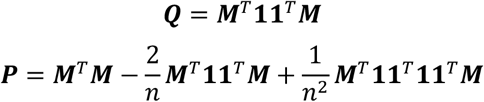 *M* denotes the community composition matrix with *m* rows of samples and *n* columns of species, where the element in i-th row and j-th column *M_ij_* is the relative abundance of species j in sample i. **1** is a unit vector with length *m* whose elements are all ones. In addition, two simple linear constrains are required to avoid ending up with an empty group or a group with all taxa that is numerically stable but ecologically trivial (Supplementary Notes).
b. continuous phenotypic variable

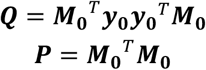 ***M*_0_** is the centered community matrix ***M*** whose column means are zero (i.e., relative abundance of each taxon is scaled by subtracting the mean of that taxon). The phenotypic vector *y* denotes that the level of phenotype at each sample is also centered to y_0_ but subtracting the mean.
c. categorical phenotypic variable

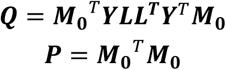 ***Y*** is an augmented categorical matrix with *m* rows and *c* columns. The row number *m* is the same as the number of total samples in the microbiome and the column number *c* is the same as the number of categories. ***L*** is a diagonal matrix whose diagonal elements are given by the inverse square root of the number of samples in each category. Optimization of the Ensemble Quotient can be achieved either by reformulating into an equivalent mixed integer linear programming problem (for small-scale problems) or by Markov-chain based heuristics like genetic algorithms (for large-scale problems). Full details of algorithms are in Supplementary Notes. R Scripts are available at Github https://github.com/Xiaoyu2425/Ensemble-Quotient-Optimization.git. An R package is currently under active development and will be posted on the same site.

### Cross-validation and aggregation network

Cross-validation consists of the following four steps. (a) Randomly splitting the dataset into a training subset and a test subset (e.g. 50-50 with the TARA oceans data). (b) Finding the best group with EQO on the training subset generated from each time of splitting. We regularized the model by finding the group that minimizes the AIC (Fig. S2A). (c) Evaluating the performance of the best assemblage generated from the training subset by calculating the cross-validation R^2^ with the corresponding test subset in each random splitting. (d) Computing the importance each single taxon as cumulative cross-validation R^2^ for assemblages where that taxon is present

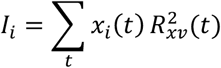

and importance of taxa pairs as cumulative cross-validation R^2^ for assemblages where the two taxa are co-present

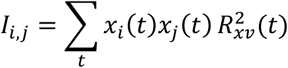

In the above expressions, *t* is the number of splitting (1~100). 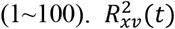 is the cross-validation R^2^ against nitrate for the test subset in the t-th splitting. *x_j_*(*t*) is 0/1 number denoting whether species *i* is present (*x_j_* = 1) or absent (*x_j_* = 0) in the group generated from the training subset in the *t*-th splitting and *x_j_*(*t*) is also a 0/1 number denoting whether species *j* is present (*x_j_* = 1) or absent (*x_j_* = 0) in the group generated from the training subset in the t-th splitting. The importance calculated above are normalized to relative importance by being divided by the maximal importance. A single taxon with high relative importance implies that this taxon tends to be included in a group with high cross-validation R^2^, while a pair of taxa with relative importance implies that those two taxa tend to be co-selected in a group with high cross-validation R^2^. The relative importance of single taxon and relative importance of taxa pairs are reflected by the size of nodes and the width of edges in an aggregation network. The aggregation network in Fig. 3B showed 11 taxa with relative importance larger than 0.5.

### Predicting gut metabolites with minimal microbiome assemblages

We ran EQO for each of the 74 metabolites that are detected in > 80% samples, for which the best group size was determined by minimizing AIC value as previously detailed. To assess the significance of metabolite prediction with minimal assemblages, we generated null expectation of linear regression R^2^ by calculating random groups with the same size for 999 times for each metabolite. A significance value (*P*-value) was evaluated from a Gaussian probability density function with the mean and standard deviation inferred from the linear regression R^2^ of random groups. Then we implemented stringent multiple hypotheses testing correction by multiplying the calculated *P*-value by a factor 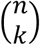 to account for the fact that the number of hypotheses to test increases rapidly with group size, where *n* is the total number of taxa in the microbiome and k is the number of taxa in the minimal assemblage. For metabolites whose minimal assemblage prediction has an adjusted *P*-value smaller than 0.01, we applied a second filter by only keeping metabolites whose minimal assemblage predictions have a cross-validated R^2^ higher than 0.2.

### Other statistical analysis

All the other statistical analyses were performed in R version 4.1.3^37^. Aggregation network visualization was performed with R package visNetwork version 2.1.0. Solution of the reformulated mixed integer programming problems were performed by Gurobi optimizer (https://www.gurobi.com) version 8.1.1 (MIT licensed) with an R 3.5.3 interface on a high-performance computing cluster at MIT. Genetic algorithm was implemented by R package GA^38,39^ version 3.2.2, with parallel computing enabled by R package doParallel 1.0.17. Fast numerical multiplication of matrices was executed by a customized C++ script, which was integrated into R script by R packages Rcpp 1.0.8 and RcppEigen 0.3.3. Visualization of other plots are performed by R package ggplot2 version 3.3.5.

## Acknowledgement

We thank Stilianos Louca, Joshua Goldford, Itzik Mizrahi and coworkers for sharing their microbiome datasets. We thank Mikhail Tikhonov and all Cordero lab members for constructive suggestions. X.S. thanks Ben Liu and Qingyi Wang for helpful discussions regarding mathematics. X.S. and O.X.C. were supported by a seed grant from the Abdul Latif Jameel Water & Food Systems Lab. O.X.C was supported by the Simons Collaboration: Principles of Microbial Ecosystems, award number 542395. A.G. is supported by the Gordon and Betty Moore Foundation as a Physics of Living Systems Fellow through grant number GBMF4513. R.G. was supported by Simons Foundation Postdoctoral Fellowship Award 653410.

## References

1. Louca, S. et al. Function and functional redundancy in microbial systems. Nature Ecology and Evolution (2018). doi:10.1038/s41559-018-0519-1

2. Anantharaman, K. et al. Thousands of microbial genomes shed light on interconnected biogeochemical processes in an aquifer system. Nat. Commun. (2016). doi:10.1038/ncomms13219

3. Louca, S., Parfrey, L. W. & Doebeli, M. Decoupling function and taxonomy in the global ocean microbiome. Science (80-.). (2016). doi:10.1126/science.aaf4507

4. Goldford, J. E. et al. Emergent simplicity in microbial community assembly. Science (80-.). 361, 469–474 (2018).

5. Nelson, M. B., Martiny, A. C. & Martiny, J. B. H. Global biogeography of microbial nitrogencycling traits in soil. Proc. Natl. Acad. Sci. U. S. A. (2016). doi:10.1073/pnas.1601070113

6. Díaz, S. & Cabido, M. Vive la différence: plant functional diversity matters to ecosystem processes. Trends Ecol. Evol. 16, 646–655 (2001).

7. Benoit, D. M., Jackson, D. A. & Chu, C. Partitioning fish communities into guilds for ecological analyses: an overview of current approaches and future directions. https://doi.org/10.1139/cjfas-2020-0455 78, 984–993 (2021).

8. Blaum, N., Mosner, E., Schwager, M. & Jeltsch, F. How functional is functional? Ecological groupings in terrestrial animal ecology: Towards an animal functional type approach. Biodivers. Conserv. 20, 2333–2345 (2011).

9. Hussain, F. A. et al. Rapid evolutionary turnover of mobile genetic elements drives bacterial resistance to phages. Science (80-.). 374, 488–492 (2021).

10. Goyal, A., Bittleston, L. S., Leventhal, G. E., Lu, L. & Cordero, O. X. Interactions between strains govern the eco-evolutionary dynamics of microbial communities. doi:10.7554/eLife.74987

11. Schindler, D. E., Armstrong, J. B. & Reed, T. E. The portfolio concept in ecology and evolution. Front. Ecol. Environ. 13, 257–263 (2015).

12. Levin, S. A. Encyclopedia of Biodiversity. (2nd ed.). (2013).

13. Shi, L. D. et al. Methane-dependent selenate reduction by a bacterial consortium. ISME J. 2021 1512 15, 3683–3692 (2021).

14. Lobb, B., Tremblay, B. J. M., Moreno-Hagelsieb, G. & Doxey, A. C. An assessment of genome annotation coverage across the bacterial tree of life. Microb. Genomics 6, (2020).

15. Schnoes, A. M., Brown, S. D., Dodevski, I. & Babbitt, P. C. Annotation Error in Public Databases: Misannotation of Molecular Function in Enzyme Superfamilies. PLOS Comput. Biol. 5, e1000605 (2009).

16. Griesemer, M., Kimbrel, J. A., Zhou, C. E., Navid, A. & D’Haeseleer, P. Combining multiple functional annotation tools increases coverage of metabolic annotation. BMC Genomics 19, 1–11 (2018).

17. Lee, D. J., Minchin, S. D. & Busby, S. J. W. Activating Transcription in Bacteria. Annu. Rev. Microbiol 66, 125–152 (2012).

18. Aidelberg, G. et al. Hierarchy of non-glucose sugars in Escherichia coli. BMC Syst. Biol. 8, 1–12 (2014).

19. Sunagawa, S. et al. Structure and function of the global ocean microbiome. Science (80-.). (2015). doi:10.1126/science.1261359

20. Gregor, R. et al. Mammalian gut metabolomes mirror microbiome composition and host phylogeny. ISME J. 2021 165 16, 1262–1274 (2021).

21. Qin, W. et al. Nitrosopumilus maritimus gen. nov., sp. nov., Nitrosopumilus cobalaminigenes sp. nov., Nitrosopumilus oxyclinae sp. nov., and Nitrosopumilus ureiphilus sp. nov., four marine ammoniaoxidizing archaea of the phylum thaumarchaeo. Int. J. Syst. Evol. Microbiol. 67, 5067–5079 (2017).

22. Schmid, M. C. et al. Anaerobic ammonium-oxidizing bacteria in marine environments: widespread occurrence but low diversity. Environ. Microbiol. 9, 1476–1484 (2007).

23. Kibe, R. et al. Upregulation of colonic luminal polyamines produced by intestinal microbiota delays senescence in mice. Sci. Reports 2014 41 4, 1–11 (2014).

24. Matsumoto, M., Kurihara, S., Kibe, R., Ashida, H. & Benno, Y. Longevity in mice is promoted by probiotic-induced suppression of colonic senescence dependent on upregulation of gut bacterial polyamine production. PLoS One 6, (2011).

25. Gupta, V. K. et al. Restoring polyamines protects from age-induced memory impairment in an autophagy-dependent manner. Nat. Neurosci. 2013 1610 16, 1453–1460 (2013).

26. Zhang, M. et al. Spermine Inhibits Proinflammatory Cytokine Synthesis in Human Mononuclear Cells: A Counterregulatory Mechanism that Restrains the Immune Response. J. Exp. Med. 185, 1759–1768 (1997).

27. Gerner, E. W. & Meyskens, F. L. Polyamines and cancer: old molecules, new understanding. Nat. Rev. Cancer 2004 410 4, 781–792 (2004).

28. Noack, J., Kleessen, B., Proll, J., Dongowski, G. & Blaut, M. Dietary Guar Gum and Pectin Stimulate Intestinal Microbial Polyamine Synthesis in Rats. J. Nutr. 128, 1385–1391 (1998).

29. Noack, J., Dongowski, G., Hartmann, L. & Blaut, M. The Human Gut Bacteria Bacteroides thetaiotaomicron and Fusobacterium varium Produce Putrescine and Spermidine in Cecum of Pectin-Fed Gnotobiotic Rats. J. Nutr. 130, 1225–1231 (2000).

30. Nakamura, A., Ooga, T. & Matsumoto, M. Intestinal luminal putrescine is produced by collective biosynthetic pathways of the commensal microbiome. Gut Microbes 10, 159–171 (2019).

31. Kitada, Y. et al. Bioactive polyamine production by a novel hybrid system comprising multiple indigenous gut bacterial strategies. Sci. Adv. 4, 62–89 (2018).

32. Datta, M. S., Sliwerska, E., Gore, J., Polz, M. F. & Cordero, O. X. Microbial interactions lead to rapid micro-scale successions on model marine particles. Nat. Commun. 7, (2016).

33. Szabo, R. E. et al. Historical contingencies and phage induction diversify bacterioplankton communities at the microscale. Proc. Natl. Acad. Sci. 119, e2117748119 (2022).

34. Yilmaz, P. et al. The SILVA and “All-species Living Tree Project (LTP)” taxonomic frameworks. Nucleic Acids Res. 42, D643–D648 (2014).

35. Quast, C. et al. The SILVA ribosomal RNA gene database project: improved data processing and web-based tools. Nucleic Acids Res. 41, D590–D596 (2013).

36. Rognes, T., Flouri, T., Nichols, B., Quince, C. & Mahé, F. VSEARCH: A versatile open source tool for metagenomics. PeerJ (2016). doi:10.7717/peerj.2584

37. R Development Core Team, R. R: A Language and Environment for Statistical Computing. R Foundation for Statistical Computing (2011). doi:10.1007/978-3-540-74686-7

38. Scrucca, L. GA: A Package for Genetic Algorithms in R. J. Stat. Softw. 53, 1–37 (2013).

39. Scrucca, L. On some extensions to GA package: hybrid optimisation, parallelisation and islands evolution. R J. 9, 187–206 (2016).

40. Gaur, A. & Arora, S. R. Solving A Quadratic Fractional Integer Programming Problem Using Linearization. Manag. Sci. Financ. Eng. 14, 25–44 (2008).

41. Yue, D., Guillén-Gosálbez, G. & You, F. Global optimization of large-scale mixed-integer linear fractional programming problems: A reformulation-linearization method and process scheduling applications. AIChE J. 59, 4255–4272 (2013).

