## Supplementary file for "Annotation-free discovery of functional groups in microbial communities"

Xiaoyu Shan[1], Akshit Goyal[2], Rachel Gregor[1], Otto X. Cordero[1*]
1 Department of Civil and Environmental Engineering, Massachusetts Institute of Technology
2 Physics of Living Systems, Department of Physics, Massachusetts Institute of Technology

The PDF file includes

**Supplementary Notes**
**1. Formulation of Ensemble Quotient**
1.1 continuous phenotypic variable
1.2 uniform phenotypic variable
1.3 categorical phenotypic variable
**2. Optimization of Ensemble Quotient**
2.1 Reformulation into a mixed integer linear programming problem (MILP)
2.2 Genetic algorithm and Aggregation Network
2.3 Boolean least square regression for a continuous phenotypic variable
**3. Interpretation of Ensemble Quotient**

**Supplementary figures S1 to S4**

**Supplementary Notes**

**1. Formulation of Ensemble Quotient**

In this section, we will prove that microbiome coarse-graining guided by a uniform phenotypic variable, a continuous phenotypic variable or a categorical phenotypic variable (Figure S1) can be generalized into a unified and simple mathematical framework featuring Ensembled Quotient

$$Ensemble\ Quotient \coloneqq \frac{\boldsymbol{x}^T\boldsymbol{Q}\boldsymbol{x}}{\boldsymbol{x}^T\boldsymbol{P}\boldsymbol{x}}, \boldsymbol{x} \in (0,1)^n$$

where $n$ is the dimension of the microbiome, e.g., the number of OTUs in a microbiome. The $\boldsymbol{x}$ is a Boolean vector of length $n$, for which 1 or 0 represent presence or absence of a species in the ensemble. For instance, $\boldsymbol{x} = [1,1,0,1,0]$ indicates that there are in total 5 species in the microbiome, in which the 1st, 2nd and 4th species are coarse-grained into the ensemble. Clearly, Ensemble Quotient has a quadratic fractional form.

**1.1 Continuous phenotypic variable**

For a continuous phenotypic variable, we need to determine which individuals should be taken into the ensemble and which ones not, so that the ensemble is strongly correlated with the external phenotypic variable of interest.

Let us start by considering a microbiome matrix $\boldsymbol{M}$ with $m$ rows of samples and $n$ columns of species, where the element in i-th row and j-th column $M_{ij}$ is the relative abundance of species j in sample i. Then the product of $\boldsymbol{M}$ and $\boldsymbol{x}$

$$\boldsymbol{s} = \boldsymbol{M}\boldsymbol{x}$$

gives us a new vector $\boldsymbol{s}$ of length $m$ whose k-th element is the relative abundance of the assemblage in the k-th sample. As for the external phenotypic variable (e.g., concentration of a metabolite), let us denote it as a vector $\boldsymbol{y}$ of length $m$, so that the l-th element of $\boldsymbol{y}$ becomes the concentration readout in sample #l. We then need to examine the correlation between the assemblage vector $\boldsymbol{s}$ and the external variable $\boldsymbol{y}$. Note that vectors $\boldsymbol{s}$, $\boldsymbol{y}$, and all the column vectors in matrix $\boldsymbol{M}$ representing each species can be scaled by subtracting their means without affecting correlations. Let us denote the scaled microbiome matrix, assemblage vector and phenotypic vector as $\boldsymbol{M_0}$, $\boldsymbol{s_0}$ and $\boldsymbol{y_0}$, respectively, since the scaling can greatly simplify the algebra below. However, note that for a uniform phenotypic variable to be discussed in section 1.3, vector scaling will not be allowed as a zero in the denominator will ruin the analytics.

Still, we have

$$\boldsymbol{s_0} = \boldsymbol{M_0}\boldsymbol{x}\ (Eq.S1)$$

For linear regression with continuous variables, the goodness of prediction is often quantified as coefficient of determination, which is statistically defined as the ratio of explained sum of squares (ESS) over the total sum of squares (TSS)

$$R^2 = \frac{ESS}{TSS} = \frac{(\widehat{\boldsymbol{y_0}} - \overline{\boldsymbol{y_0}})^T(\widehat{\boldsymbol{y_0}} - \overline{\boldsymbol{y_0}})}{(\boldsymbol{y_0} - \overline{\boldsymbol{y_0}})^T(\boldsymbol{y_0} - \overline{\boldsymbol{y_0}})}$$

For scaled $\boldsymbol{y_0}$ we have the mean $\overline{\boldsymbol{y_0}} = \boldsymbol{0}$. Combining the projection formulation in linear algebra

$$\widehat{\boldsymbol{y_0}} = \frac{\boldsymbol{s_0}^T\boldsymbol{y_0}}{\boldsymbol{s_0}^T\boldsymbol{s_0}}\boldsymbol{s_0}$$

we have

$$R^2 = \frac{\boldsymbol{s_0}^T\boldsymbol{y_0}\boldsymbol{y_0}^T\boldsymbol{s_0}}{\boldsymbol{s_0}^T\boldsymbol{s_0} \cdot \boldsymbol{y_0}^T\boldsymbol{y_0}}$$

in which $\boldsymbol{y_0}^T\boldsymbol{y_0}$ can be neglected in the optimization since it is just comprised of known constants. Plugging in (Eq. S1), the problem of maximizing $R^2$ can be reformulated as the following integer programming with an objective function in the form of quadratic quotient

$$max\ \frac{\boldsymbol{x}^T\boldsymbol{Q}\boldsymbol{x}}{\boldsymbol{x}^T\boldsymbol{P}\boldsymbol{x}}$$

$$s.t.\boldsymbol{x} \in (0,1)^n$$

where

$$\boldsymbol{Q} = \boldsymbol{M_0}^T\boldsymbol{y_0}\boldsymbol{y_0}^T\boldsymbol{M_0}$$

$$\boldsymbol{P} = \boldsymbol{M_0}^T\boldsymbol{M_0}$$

Note that the coefficient of determination $(R^2)$ for linear regression is numerically equivalent to Pearson's correlation coefficient in its square form $(r^2)$, although they are based on slightly different statistical contexts. In theory, it is possible that maximizing $R^2$ might lead to an assemblage with very strong negative correlation with the phenotype. This is in fact not an issue for microbiome datasets due to its compositionality, since the remaining set of species will automatically have strong positive correlation with the phenotype (same correlation strength but just with the opposite sign). In a few cases where the sign of the optimized assemblage indeed matters, we can adopt a modified formulation to maximize Pearson's correlation coefficient, but not in its square form

$$r^2 = \frac{Cov(\boldsymbol{s_0},\boldsymbol{y_0})}{\sigma(\boldsymbol{s_0})\sigma(\boldsymbol{y_0})} = \frac{\boldsymbol{s_0}^T\boldsymbol{y_0}}{\sqrt{\boldsymbol{s_0}^T\boldsymbol{s_0}\cdot\boldsymbol{y_0}^T\boldsymbol{y_0}}}$$

which can be also rewritten with $\boldsymbol{x}$ as the unknown variable as

$$max\frac{\boldsymbol{x}^T\boldsymbol{M_0}^T\boldsymbol{y_0}}{\sqrt{\boldsymbol{x}^T\boldsymbol{M_0}^T\boldsymbol{M_0}\boldsymbol{x}}}$$

**1.2 Categorical phenotypic variable**

The case with categorical phenotypic variable (healthy vs. disease, or several different sub-types of diseases) is conceptually the same as the case with continuous phenotypic variable. For a categorical phenotypic variable, an equivalent statistical metric to the coefficient of determination can be also constructed by the sum of square between/among treatments against the total of sum of square

$$R^2 = \frac{SS_{bewteen\ treatment}}{SS_{total}} = \frac{(\boldsymbol{s_0}^T\boldsymbol{Y}\boldsymbol{L})^2}{\boldsymbol{s_0}^T\boldsymbol{s_0}}$$

where $\boldsymbol{Y}$ is an augmented categorical matrix with $m$ rows and $c$ columns. The row number $m$ is the same as the number of total samples and the column number $c$ is the same as the number of categories. For instance, if there are altogether 5 samples in the microbiome dataset where samples 1, 2, 4 are from healthy hosts and samples 3,5 from hosts with a disease, then the matrix $\boldsymbol{Y}$ will be

$$\boldsymbol{Y} = \begin{bmatrix} 1 & 0 \\ 1 & 0 \\ 0 & 1 \\ 1 & 0 \\ 0 & 1 \end{bmatrix}$$

$\boldsymbol{L}$ is a diagonal matrix whose diagonal elements are given by the inverse square root of the number of samples in each category.

Combining (Eq. S1) again, we get

$$R^2 = \frac{\boldsymbol{x}^T \boldsymbol{M_0}^T \boldsymbol{YLL}^T \boldsymbol{Y}^T \boldsymbol{M_0 x}}{\boldsymbol{x}^T \boldsymbol{M_0}^T \boldsymbol{M_0 x}}$$

Clearly, we again reach the same quadratic quotient form of

$$min \frac{\boldsymbol{x}^T \boldsymbol{Qx}}{\boldsymbol{x}^T \boldsymbol{Px}}$$

$$s.t. \boldsymbol{x} \in (0,1)^n$$

where

$$\boldsymbol{Q} = \boldsymbol{M_0}^T \boldsymbol{YLL}^T \boldsymbol{Y}^T \boldsymbol{M_0}$$

$$\boldsymbol{P} = \boldsymbol{M_0}^T \boldsymbol{M_0}$$

**1.3 Uniform phenotypic variable**

With a uniform phenotypic variable, we are not allowed to directly construct a correlation/regression-type of $R^2$ although in spirit it can be also regarded as special correlation/regression problem with a constant number across all samples. Doing so implies that the variance of the phenotypic variable is zero, which becomes illegal as a denominator.

Here, we examine the coefficient of variation of the ensemble, which is commonly adopted to quantify the level of variation across samples. In statistics, the coefficient of variation is defined as standard deviation divided by mean,

$$CV = \frac{\sigma}{\mu} \; (Eq. S2)$$

As aforementioned, here we are no longer able to simply the expression of variance by scaling the vectors as $\mu(\boldsymbol{s}) = 0$ is not acceptable as a denominator. To derive the full expression of the variance of $\boldsymbol{s}$, we need an auxiliary unit vector $\boldsymbol{1}$ with length $m$ whose elements are all ones. Then we have

$$nVar(\boldsymbol{s}) = (\boldsymbol{s} - \mu\boldsymbol{1})^T(\boldsymbol{s} - \mu\boldsymbol{1}) \; (Eq. S3)$$

Where $\mu$, a scalar, denotes the mean of the ensemble $\boldsymbol{s}$, which can be at the same time expressed as

$$n \cdot \mu = \boldsymbol{1}^T\boldsymbol{s} = \boldsymbol{s}^T\boldsymbol{1} \; (Eq. S4)$$

Combining (Eq. S1~S4), we can derive that

$$CV^2 \sim \frac{\boldsymbol{x}^T \left(\boldsymbol{M}^T\boldsymbol{M} - \frac{2}{n}\boldsymbol{M}^T\boldsymbol{11}^T\boldsymbol{M} + \frac{1}{n^2}\boldsymbol{M}^T\boldsymbol{11}^T\boldsymbol{11}^T\boldsymbol{M}\right)\boldsymbol{x}}{\boldsymbol{x}^T(\boldsymbol{M}^T\boldsymbol{11}^T\boldsymbol{M})\boldsymbol{x}}$$

Again, we arrive at the form of the microbiome ensemble quotient. Similarly, we are able to reformulate coarse-graining problem into the following integer programming format

$$max \frac{\boldsymbol{x}^T \boldsymbol{Qx}}{\boldsymbol{x}^T \boldsymbol{Px}}$$

$$s.t. \boldsymbol{x} \in (0,1)^n$$

where

$$\boldsymbol{Q} = \boldsymbol{M}^T\boldsymbol{11}^T\boldsymbol{M}$$

$$\boldsymbol{P} = \boldsymbol{M}^T\boldsymbol{M} - \frac{2}{n}\boldsymbol{M}^T\boldsymbol{11}^T\boldsymbol{M} + \frac{1}{n^2}\boldsymbol{M}^T\boldsymbol{11}^T\boldsymbol{11}^T\boldsymbol{M}$$

Conceptually, the efforts to minimize the coefficient of variation is analogous to maximize the “correlation” with a uniform vector (e.g., $\boldsymbol{y} = [1,1,1,1,1]$) although it is illegal in practice since such uniform vectors have zero variance. Nevertheless, this analogy can be helpful for understanding the generalization of stabilizing grouping and associating grouping into the same formulation.

In addition, two simple linear constrains are required to avoid generating an empty group or a group with all taxa that are numerically stable but ecologically trivial

$$(\boldsymbol{M}^T \mathbf{1}_m)^T \boldsymbol{x} \geq me$$
$$(\boldsymbol{M}^T \mathbf{1}_m)^T \boldsymbol{x} \leq m(1-e)$$

In those constraints, $\mathbf{1}_m$ is a unit vector with length $m$ whose elements are all ones. These constraints imply that the average abundance of the aggregated group across all the samples should be no smaller than a threshold given by $e$ (for instance 5%) and no larger than a threshold given by $1-e$ (for instance 95%).

### 2. Optimization of Ensemble Quotient

#### 2.1 Reformulation into a mixed integer linear programming (MILP) problem

The binary nature of coarse-graining provides the ensemble quotient with inherent mathematical simplicity. It has been shown in integer programming studies that quadratic fractional optimization, when the variables are integers, can be reduced to an equivalent mixed integer linear programming problem[40]. The method includes Charnes-Cooper transformation incorporated with Glover’s linearization, which has been widely used in solving fractional programming problems[41]. Briefly, the reformulation-linearization algorithm works by introducing new variables with linear constraints. In this sense, the above quadratic fractional programming problem can be reformulated into

$$min\ \mathbf{1}^T \boldsymbol{\alpha} - m_p \boldsymbol{k}$$

$$s.t.\ \boldsymbol{\alpha} + \boldsymbol{\beta} - \boldsymbol{P}\boldsymbol{k} - m_p u \mathbf{1} = \mathbf{0}\ (C1)$$
$$\boldsymbol{\alpha} - 2m_p u \mathbf{1} + 2m_p \boldsymbol{k} \leq \mathbf{0}\ (C2)$$
$$\boldsymbol{\beta} - 2m_p \boldsymbol{k} \leq \mathbf{0}\ (C3)$$
$$\boldsymbol{\gamma} + \boldsymbol{\delta} - \boldsymbol{Q}\boldsymbol{k} - m_q u \mathbf{1} = \mathbf{0}\ (C4)$$
$$\boldsymbol{\gamma} - 2m_q u \mathbf{1} + 2m_q \boldsymbol{k} \leq \mathbf{0}\ (C5)$$
$$\boldsymbol{\delta} - 2m_q \boldsymbol{k} \leq \mathbf{0}\ (C6)$$
$$\mathbf{1}^T \boldsymbol{\delta} - m_q \boldsymbol{k} = \mathbf{1}\ (C7)$$
$$\boldsymbol{k} - u\mathbf{1} - R\boldsymbol{x} \geq -R\mathbf{1}\ (C8)$$
$$\boldsymbol{k} - u\mathbf{1} \leq \mathbf{0}\ (C9)$$
$$\boldsymbol{k} - R\boldsymbol{x} \leq \mathbf{0}\ (C10)$$
$$u - R \leq 0\ (C11)$$

Where $\boldsymbol{\alpha}, \boldsymbol{\beta}, \boldsymbol{\gamma}, \boldsymbol{\delta}, \boldsymbol{k} \geq \mathbf{0}$ are vectors of length $n$, $\boldsymbol{x} \in (0,1)^n$ is the binary variable to be solved through optimization, $R$ is a sufficiently large number, $m_p$ and $m_q$ are defined as an upper bound of row sums of $\boldsymbol{P}$ and $\boldsymbol{Q}$, respectively.

$$m_p = \max_i\left(\sum_j |P_{ij}|\right)$$

$$m_q = \max_i\left(\sum_j |Q_{ij}|\right)$$

In this way, the binary quadratic fraction programming problem with $n$ unknown variables is reformulated into an MILP with $6n+1$ unknown variables embedded with $9n+2$ linear constraints. The reformulated MILP is directly accessible by commercially available integer optimizers such as Gurobi and Cplex. However, MILP itself is still NP-hard in nature, making this only applicable to small-scale problems.

### 2.2 Genetic algorithm and Aggregation Network

An alternative approach to optimize the Ensemble Quotient to leverage heuristic algorithms. In particular, the binary nature of the unknown variable $x$ in the formulation makes Genetic Algorithm an ideal candidate. Briefly, the presence or absence of a species in the assemblage is to be regarded as dominance or recessive of a loci in a genetic algorithm. Then a Markov-chain based stochastic search is implemented to simulate the mutation, recombination and selection of the "genotype" based on its fitness given by the object function for optimization. In our case, the fitness function is the Ensemble Quotient. In this way, the optimal assemblage given by the genetic algorithm is analogous to a genotype "evolved" towards the peak in the fitness landscape.

A potential shortcoming of genetic algorithm lies in the fact that it only converges in probability as heuristics. However, for microbiome studies we are more interested in gaining biological insights from the most robust statistical patterns than recovering the very exact optimal solutions in complex high-dimensional datasets. In light of that, we have developed a cross-validation-based algorithm to capture and characterize the most important cohesive guilds. Briefly, the relative importance of single species and species pairs are evaluated as cumulative cross-validation $R^2$ for assemblages where that single species or the species pair is present, which can be visualized as the node size and edge width in an Aggregation Network (Methods). One can then infer the most important species that should be grouped together by examining strongly connected big nodes in an Aggregation network, as in the Tara Oceans case (Figure 3B).

### 2.3 Boolean least square regression for a continuous phenotypic variable

Specifically, when the phenotypic variable is continuous, the Ensemble Quotient can be reformulated into the most ordinary least square format. In addition to the slope $k$ and intercept $b$, we also need Boolean variables to indicate the presence or absence of each species in the assemblage.

$$\boldsymbol{y} \sim k\left(\sum_j x_j \boldsymbol{m_j}\right) + b$$

Where $\boldsymbol{m_j}$ is column $j$ in microbiome matrix $\boldsymbol{M}$ indicating the relative abundance of species $j$ in each sample. Again, $x_j$, the binary variable, denotes whether species $j$ should be included in the assemblage. Then the least square error is

$$\boldsymbol{e^2} = ||\boldsymbol{Az} - \boldsymbol{y}||_2 = \boldsymbol{z}^T\boldsymbol{A}^T\boldsymbol{Az} - \boldsymbol{2y}^T\boldsymbol{Az} + \boldsymbol{y}^T\boldsymbol{y}$$

Where $\boldsymbol{A}_{m\times(n+1)}$ is the augmented microbiome matrix $[\mathbf{1}_m \vdots \boldsymbol{M}_{m\times n}]$ since we need an additional column of 1 to map the linear intercept. $\boldsymbol{z}$ is the augmented vector for unknown variable $[b \vdots k\boldsymbol{x}]$. Also noting that $\boldsymbol{y}^T\boldsymbol{y}$ can be neglected as a known term, we have

$$min\ \boldsymbol{z}^T\boldsymbol{Q}\boldsymbol{z} + \boldsymbol{L}\boldsymbol{z}$$

where $\boldsymbol{Q} = \boldsymbol{A}^T\boldsymbol{A},\ \boldsymbol{L} = -2\boldsymbol{y}^T\boldsymbol{A}$.

In this way, we are going to solve a mixed integer quadratic programming problem, which is also solvable by commercial optimizer such as Gurobi (later than v8.0).

#### 3. Interpretation of Ensemble Quotient

Here we discuss the simplest case with a continuous phenotypic variable. It can be easily generalized since categorical/uniform phenotypic variables can be regarded as special forms of continuous phenotype cases as detailed in section 1. Recall the expression of $R^2$ for a continuous phenotypic variable as

$$R^2 \sim EQ = \frac{(\boldsymbol{y_0}^T\boldsymbol{s_0})^2}{\boldsymbol{s_0}^T\boldsymbol{s_0}}$$

Which can be rewritten as

$$EQ = \frac{[\boldsymbol{y}^T(\boldsymbol{S}\cdot\mathbf{1})]^2}{\mathbf{1}^T\boldsymbol{S}^T\boldsymbol{S}\mathbf{1}}$$

Where $\boldsymbol{S}$ is the assemblage matrix (the subset of the whole microbiome matrix $\boldsymbol{M}$ but only with species included in the assemblage). Again, neglecting the constant term $\boldsymbol{y}^T\boldsymbol{y}$, the expression above can be transformed into a form that is more statistically interpretable,

$$EQ = \frac{(\overline{cov_{(\boldsymbol{x_i},\boldsymbol{y})}})^2}{\overline{cov_{(\boldsymbol{x_i},\boldsymbol{x_j})}}}$$

Where $\overline{cov_{(\boldsymbol{x_i},\boldsymbol{y})}}$ means the average covariance between each species and the phenotypic variable, $\overline{cov_{(\boldsymbol{x_i},\boldsymbol{x_j})}}$ means the average covariance between each species and each species. In the simplest scenario where all vectors have unit standard deviation, it can be further simplified as

$$EQ = \frac{(\overline{r_{(\boldsymbol{x_i},\boldsymbol{y})}})^2}{\overline{r_{(\boldsymbol{x_i},\boldsymbol{x_j})}}}$$

Which basically reflects the ratio of average species-phenotype correlation and average species-species correlation. On one hand, if species within an assemblage have opposite signs of correlation with a phenotype, then their average would be nearly zero, resulting in small value of the nominator (i.e., poor consistency). On the other hand, if species within an assemblage tend to be strongly correlated with each other, then their average would lead to a large value of the denominator (i.e., poor complementarity). An optimal assemblage should be formed by species with strong and consistent correlations with the phenotype but weak or even negative correlations with each other.

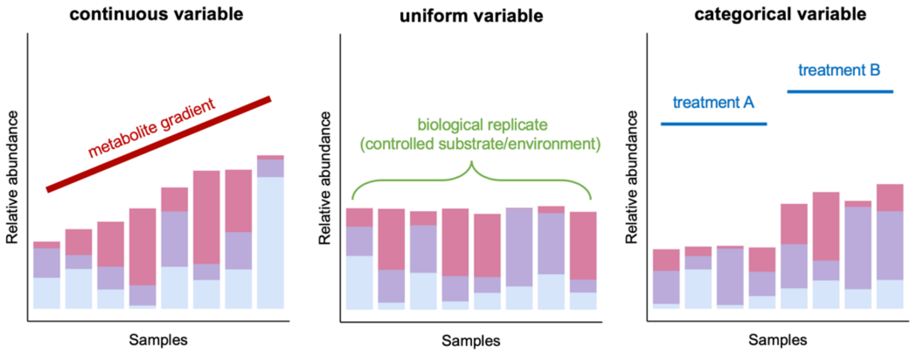


**Figure S1 Schematic illustration of different types of environmental/phenotypic variables that can be leveraged by EQO to predict functional groups**. Bar plots show relative abundance of different taxa. Appropriate grouping of taxa leads to strong coupling with the phenotypic variable. For a continuous phenotypic variable such as measured concentration of a metabolite across samples, the coupling can be captured by a strong correlation. For a uniform phenotypic variable, the coupling can be marked as high stability or low variability. For a categorical phenotypic variable, the coupling can be interpreted as significant discrimination between treatment.

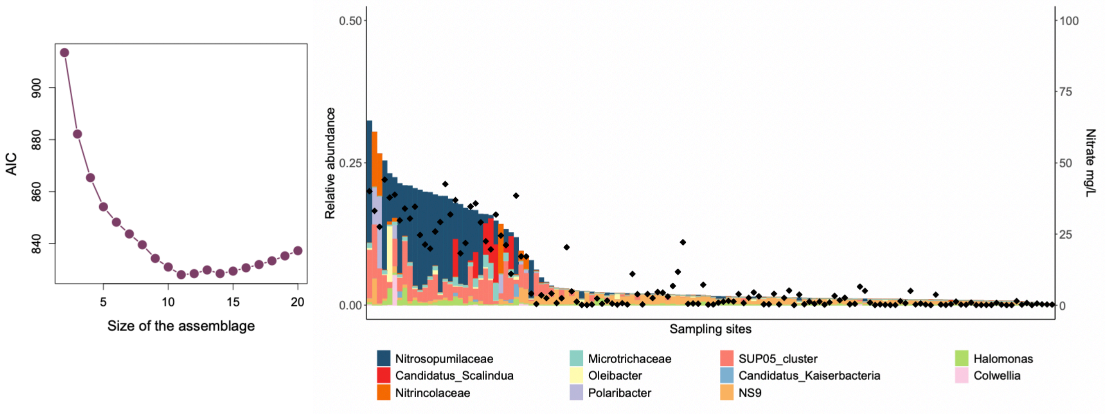


**Figure S2 EQO for Tara Ocean microbiome.** (A) The best group size for EQO was determined to be 11 based on an AIC minimization criterion. (B) Relative abundance distribution of the 11 taxa selected by the algorithm across all sampling sites (left y-axis). Nitrate concentration measured at each sampling site was shown as black dots (right y-axis).

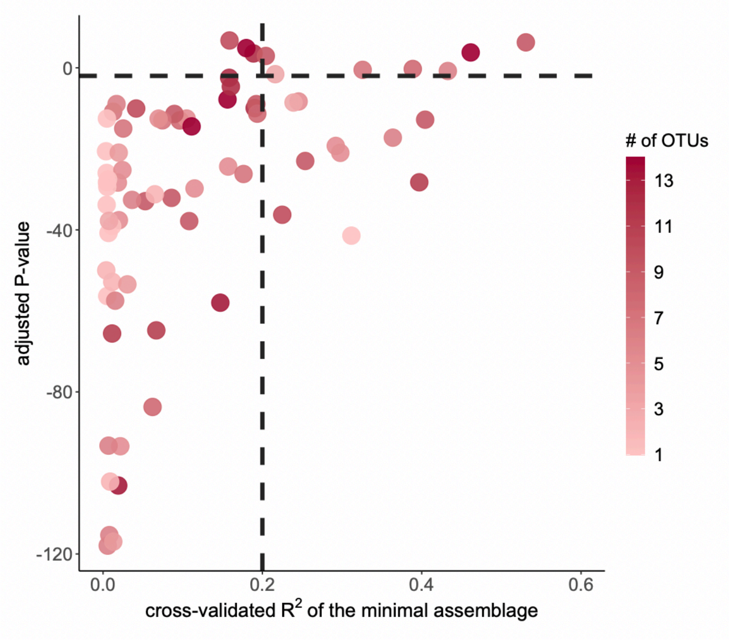


**Figure S3 Statistical tests for predictability of metabolite in the gut microbiome dataset.** Metabolites with cross-validated $R^2$ lower than 0.2 as well as adjusted P-value higher than 0.01 were filtered out from further analysis (Methods).

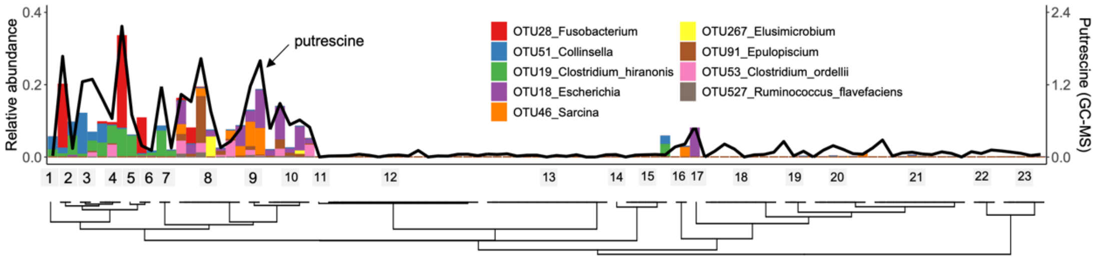


**Figure S4 Distribution of predicted putrescine functional group in samples ranked according to host phylogeny.** 1, hyena; 2, tiger; 3, leopard; 4, lion; 5, sand cat; 6, jungle cat; 7, wolf; 8, coati; 9, black bear; 10, brown bear; 11, donkey; 12, zebra; 13, rhino; 14, goat; 15, sheep; 16, lemur; 17, capuchin; 18, mandrill; 19, gibbon; 20, gorilla; 21, chimpanzee; 22, African elephant; 23, Asian elephant.
